# Mesoscale developmental rivalry in human extrastriate visual cortex

**DOI:** 10.1101/2025.10.13.682053

**Authors:** Shahin Nasr, Jan Skerswetat, Bryan Kennedy, Marianna E. Schmidt, Eric D. Gaier, Antony B. Morland, Peter Bex, David G. Hunter

## Abstract

In humans and non-human primates, the extrastriate visual cortex contains fine-scale columns selectively responsive to motion, disparity, and color. However, the developmental interplay between these functional modules remains poorly understood. Using high-resolution fMRI, we compared the mesoscale organization of extrastriate cortex in 16 individuals with normal vision and 15 participants with amblyopia (PwA) caused by strabismus (n=8) or anisometropia (n=7). In controls, the cortical territory occupied by disparity-selective columns exhibited a competitive relationship with that of motion- and color-selective columns. Consistent with this pattern, PwA showed a reduction in disparity-selective responses accompanied by enhanced motion- and color-selective activity, as well as expansion of the cortical territory allocated to them. Our results show that the mesoscale modules of the human visual system are rivals in development allowing intact functions to usurp those that are compromised.

## 1. Introduction

The human visual system operates under a constant flood of rich sensory input, with features such as color, motion, depth, and shape that must be encoded to guide behavior and enhance survival. Yet processing resources are finite, likely forcing specialized pathways for different visual features to compete. This competition emerges during early development, when neuronal functions are shaped and consolidated by visual experience. However, our understanding of how specialized neuronal functions rival during development – from here on referred to as ***developmental rivalry*** - remains limited. This is in part due to the challenges of resolving mesoscale neuronal sites dedicated to specific visual features and the impossibility of experimentally manipulating developmental trajectories in humans. Studying the developmental disorder amblyopia—caused by disrupted binocular vision— therefore offers one of a very few opportunities to probe the mechanisms underlying developmental rivalry in the human brain.

In the primary visual cortex, amblyopia caused by either strabismus or anisometropia leads to a reduction in the number of binocular neurons (Crawford and von Noorden, 1979; Movshon et al., 1987; Smith et al., 1997; Kiorpes et al., 1998; Bi et al., 2011). Studies in non-human primates (NHPs) suggest that this deficit propagates to downstream visual areas, resulting in fewer disparity-sensitive neurons in regions such as V2 and V4 (Kiorpes et al., 1998; Bi et al., 2011). However, to our knowledge, the consequences of these developmental changes for the mesoscale functional organization of the other sites across early visual areas have not yet been reported. Specifically, it is unclear whether the reduction in disparity-selective neurons triggers a competitive expansion of cortical columns in early visual areas that encode visual features that are mostly spared by amblyopia (e.g., motion and color), indicative of system-wide functional reorganization and if this phenomenon varies depending on the amblyopia cause.

In the extrastriate visual cortex of humans and NHPs, disparity-selective columns are interdigitated with color- and motion-selective columns (Nasr et al., 2016; Li et al., 2019; Kennedy et al., 2023; Wang et al., 2024). In the earlier areas such as V2, columns with alike stimulus preference are organized in systematic, parallel stripe-like patterns that originate at the V1–V2 border and extend into higher-order visual areas. While the impact of amblyopia on the mesoscopic features expressed in the ocular dominance columns in primary visual cortex have been examined in multiple studies (Crawford and von Noorden, 1979; Kiorpes et al., 1998; Goodyear et al., 2002; Nasr et al., 2025), the impact on the mesoscopic organization of extrastriate cortex, beyond reduced disparity-selective activity, has not yet been explored.

Over the past decade, high-resolution fMRI has been used to visualize the functional organization of cortical columns in the human extrastriate cortex that are selectively involved in encoding color (Nasr et al., 2016; Nasr and Tootell, 2018a; Navarro et al., 2021), motion (Tootell and Nasr, 2021; Kennedy et al., 2023) and disparity (Goncalves et al., 2015; Nasr et al., 2016; Nasr and Tootell, 2018b). Analogous results have been also reported in NHPs based on using similar techniques (Li et al., 2019; Wang et al., 2024).

In the present study, we applied this technique to investigate the mesoscopic functional organization of extrastriate visual areas V2-V4 in individuals with normal vision as well as those with strabismic or anisometropic amblyopia. We examined whether the reduced disparity-selective activity observed in the participants with strabismic vs. anisometropic amblyopia is accompanied by upregulation of motion- and color-selective responses and expansion of columns functionally selective to motion and color, compared to controls. Our results, confirming this prediction, provide one of the first lines of evidence for developmental rivalry in the mesoscale organization of extrastriate cortex of humans.

## 2. Results

Fifteen participants with amblyopia (PwA; Table S1)—including eight with strabismus (4 females) and seven with anisometropic amblyopia (3 females)—and sixteen controls (5 females) took part in the study. Age did not differ significantly between PwA and controls (t(29) = 0.09, *p* = 0.93) or between strabismic and anisometropic individuals (t(13) = 0.87, *p* = 0.40). The interocular visual acuity difference—an index of amblyopia severity—was also not statistically different between strabismic and anisometropic participants (t(13) = 1.26, *p* = 0.25). All participants had normal color perception, verified using Farnsworth D-15 test. Each participant underwent five scanning sessions on separate days to measure disparity-, motion-, and color-selective responses, define retinotopic area boundaries, and acquire structural data (see Methods).

### 2.1. Loss of disparity-selective responses accompanies poor stereovision in amblyopia

PwA showed reduced stereoacuity and this impairment was accompanied by a marked reduction in disparity-selective activity in individuals with amblyopic vision (Fig. 1). Specifically, in participants with normal vision, disparity-selective responses in V2 formed a distinct stripe-like pattern that extends into downstream visual areas along both the ventral (e.g., V4) and dorsal (e.g., intraparietal sulcus) streams (Goncalves et al., 2015; Nasr et al., 2016; Kennedy et al., 2023). These mesoscale disparity-selective sites were interdigitated with regions preferring zero-disparity stimuli. In PwA, disparity-selective sites were rarely detectable in extrastriate cortex, whereas some zero-disparity–preferring sites remained evident across visual areas.

**Figure 1).**
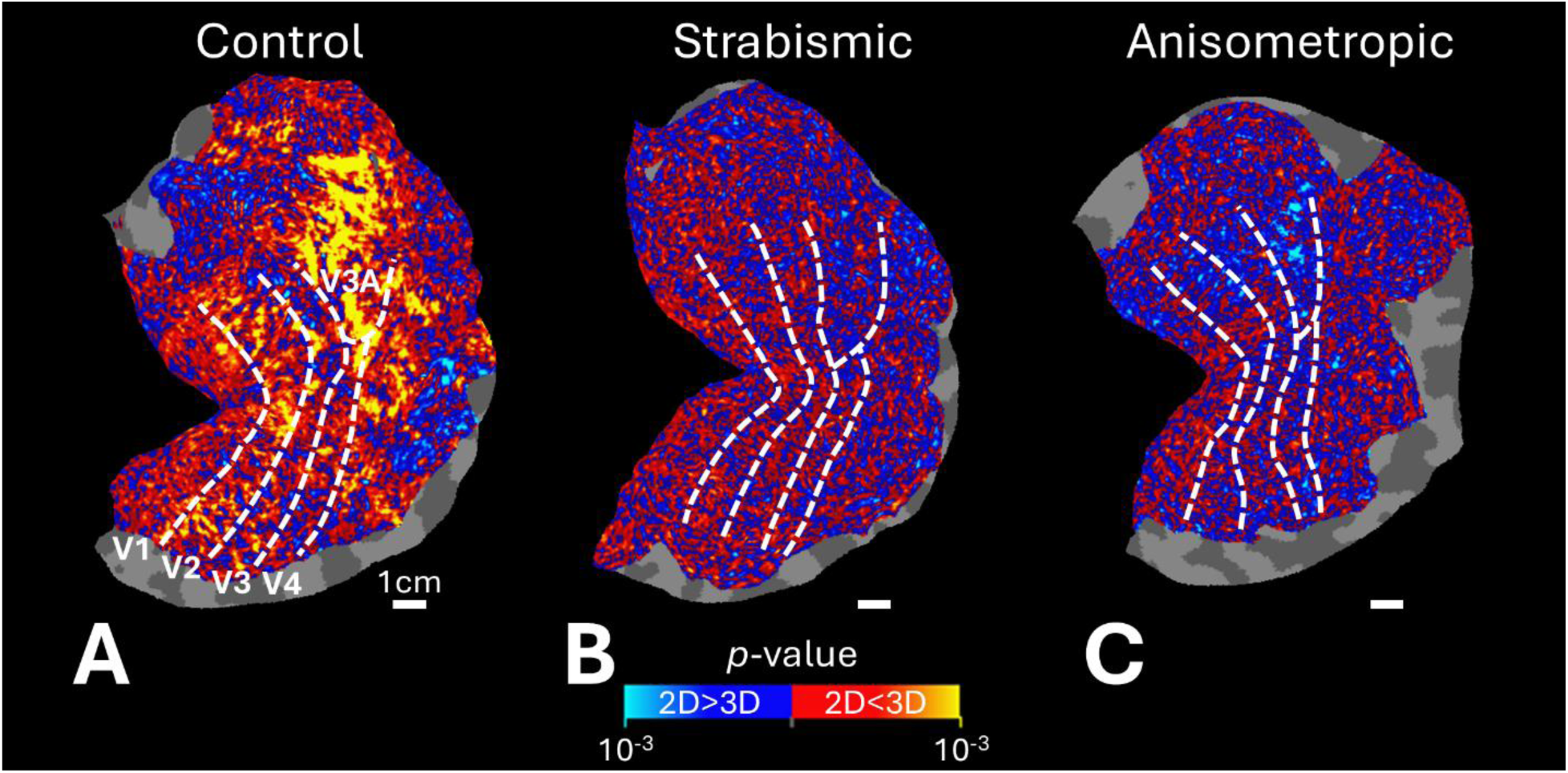
Disparity-selective activity in extrastriate visual areas of individuals with neurotypical and amblyopic vision. Panels **A–C** show unthresholded activity evoked by contrasting disparity-varying versus zero-disparity stimuli in an individual with neurotypical, strabismic, and anisometropic vision, respectively. In the neurotypical individual, disparity-selective sites form stripy patterns in V2, with activity extending into higher-level visual areas V3–V4. In the strabismic and anisometropic individuals with poor stereoacuity, disparity-selective activity is markedly reduced. White lines indicate retinotopically defined visual area borders. The white scale bar represents 1 cm.

The selective responses to disparity are significantly larger in controls than PwA across all areas (F(1,29) = 16.57, *p* < 10^-3^). This main effect was accompanied by an interaction between participant group and visual area (F(2,58) = 20.94, *p* < 10^-4^), reflecting a reduction in between-group differences higher in the visual hierarchy (Fig. 2). In contrast, there was no corresponding between-group difference in the amplitude of selective response in zero-disparity preferring sites (F(1,29) = 0.06, *p* = 0.81) and no interaction between group and visual area (F(2,58) = 2.94), p = 0.07) (Table S2). No significant difference between the two amblyopia subtypes (F(1,13) < 0.40, *p* = 0.54) was present and no interaction with visual area either (F(2,26) = 1.76, *p* = 0.20) (Table S3). Thus, PwA showed a decline in the level of disparity-selective responses, whereas activity within zero-disparity preferring sites remained comparable to controls.

**Figure 2).**
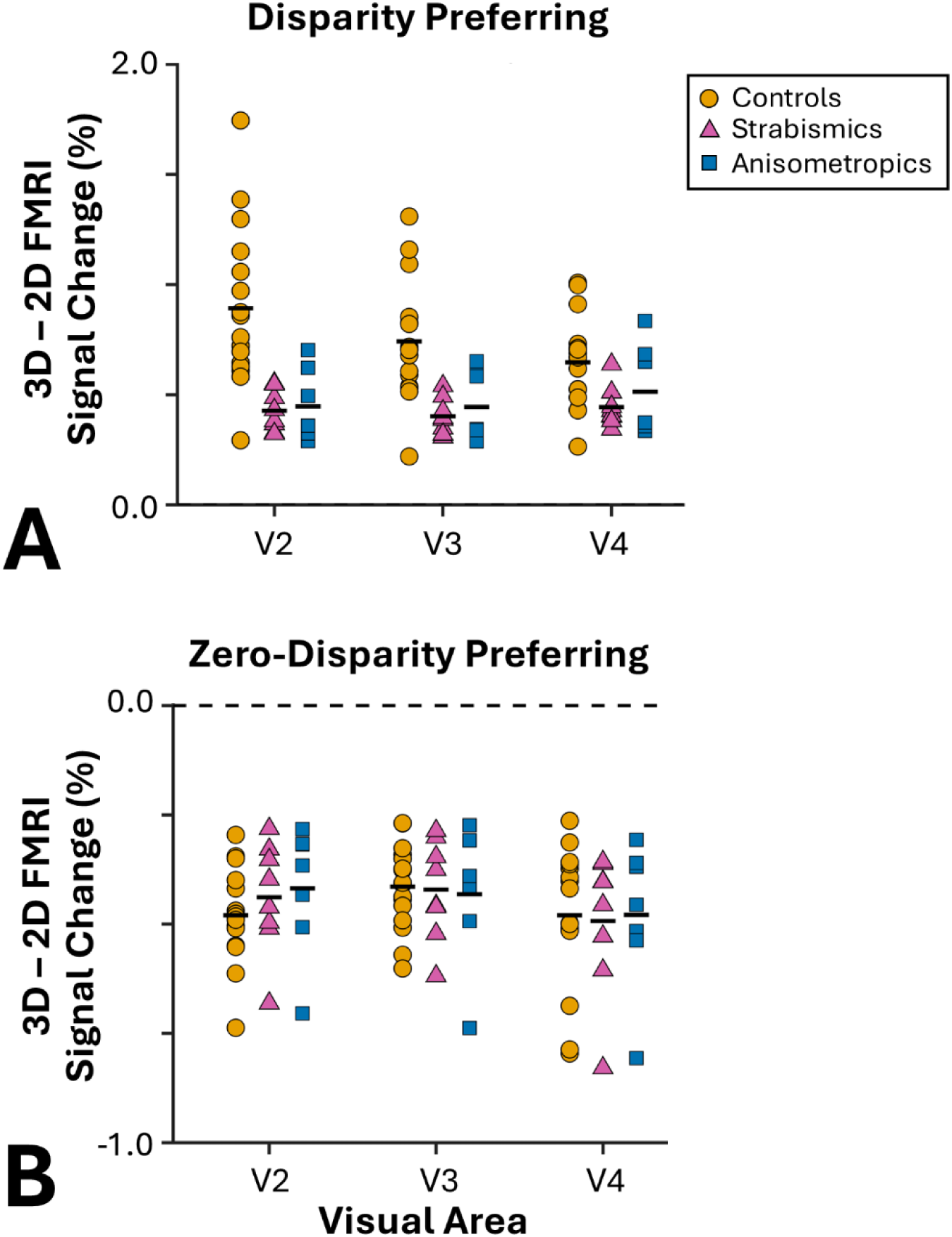
Quantified disparity-selective responses across areas V2–V4. For each participant, visual areas were divided into regions preferring disparity-varying versus zero-disparity stimuli, based on the unthresholded maps. In regions preferring disparity-varying stimuli (top panel), controls showed stronger responses compared with PwA. In regions preferring zero-disparity stimuli (bottom panel), the level of disparity-selective activity was comparable across groups. Error bars indicate ±1 S.E.M.

We next examined whether the amplitude of evoked activity could be predicted from the interocular visual acuity difference while accounting for visual area (V2, V3, V4) (Fig. 3). Application of a regression analysis to the data from PwA revealed that disparity-selective activity significantly decreased with increasing interocular visual acuity difference (R^2^ adjusted = 0.18, t = −2.13, *p* = 0.03), without any significant interaction with the effect of area (t < 0.52, *p* > 0.60). These results indicate that disparity-selective responses decline with increasing severity of amblyopia. Notably, most of our amblyopic participants did not show any measurable stereoacuity based on Randot test. In this condition, we could not check the relationship between the residual disparity-selective activity and the behaviorally measured stereoacuity.

**Figure 3).**
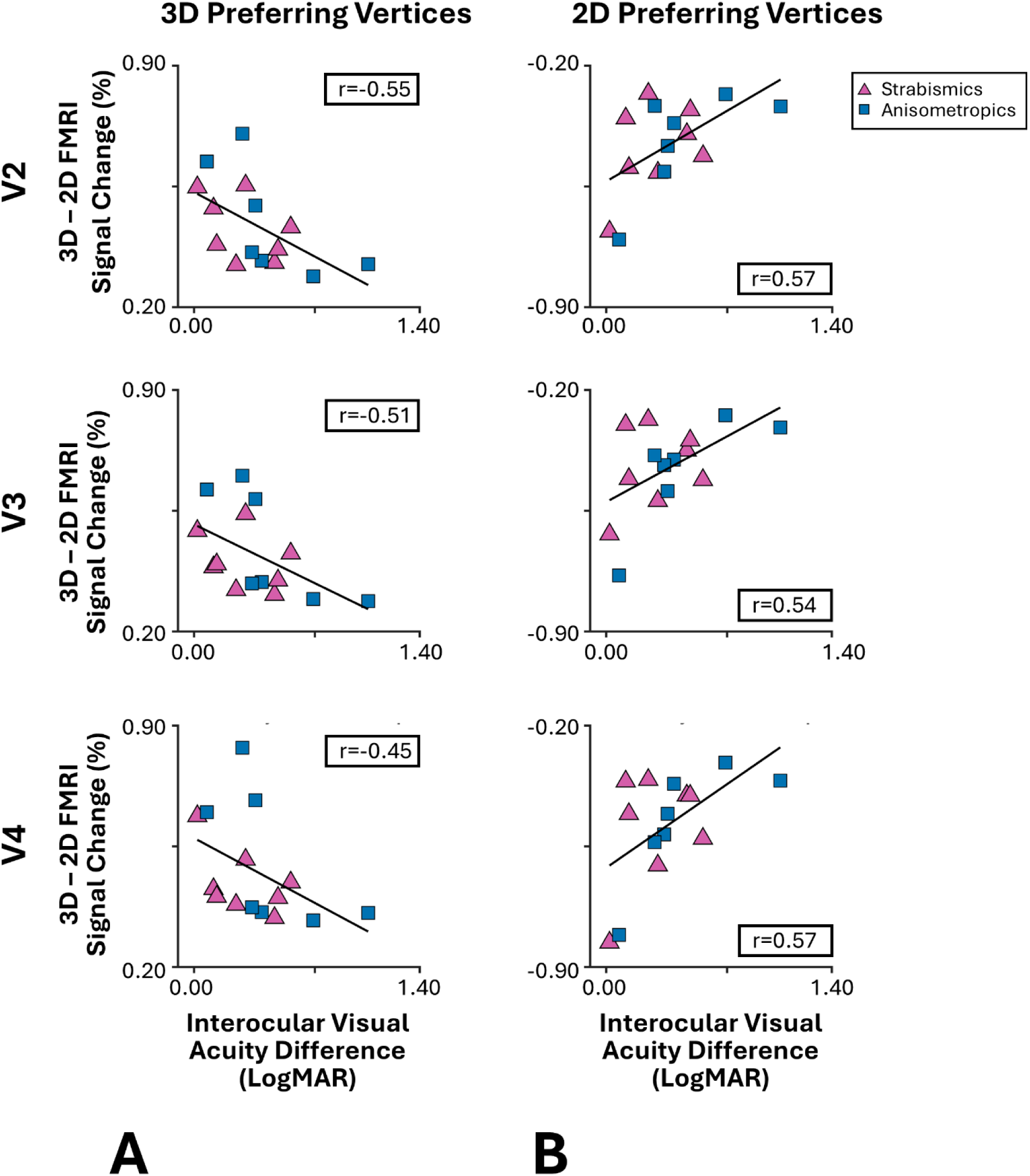
Correlation between disparity-selective activity and amblyopia severity. Across regions preferring disparity-varying stimuli (top panels), disparity-selective activity negatively correlated with interocular visual acuity difference—i.e., greater amblyopia severity was associated with weaker disparity-selective responses. In regions preferring zero-disparity stimuli (bottom panels), an opposite relationship was observed. Each data point represents one participant with either strabismic (red) or anisometropic (blue) amblyopia.

### 2.2. Increased motion- and color-selective activity in amblyopia

We hypothesized that the decrease in the level of disparity-selective activity might be associated with an increase in the level of motion- and color-selective response. To test this, we next examined the response to moving (vs. stationary) and color-varying (vs. luminance-varying) stimuli. As demonstrated in Fig. 4, the overall pattern of evoked activity, including the expected stripy patterns in areas V2-V4, remained visibly comparable across all participants. In controls, motion-, disparity- and color-selective sites form an interdigitated organization with small overlap between them. In amblyopic individuals, this interdigitated organization is mostly preserved (for color and motion), even in the absence of disparity-selective sites.

**Figure 4).**
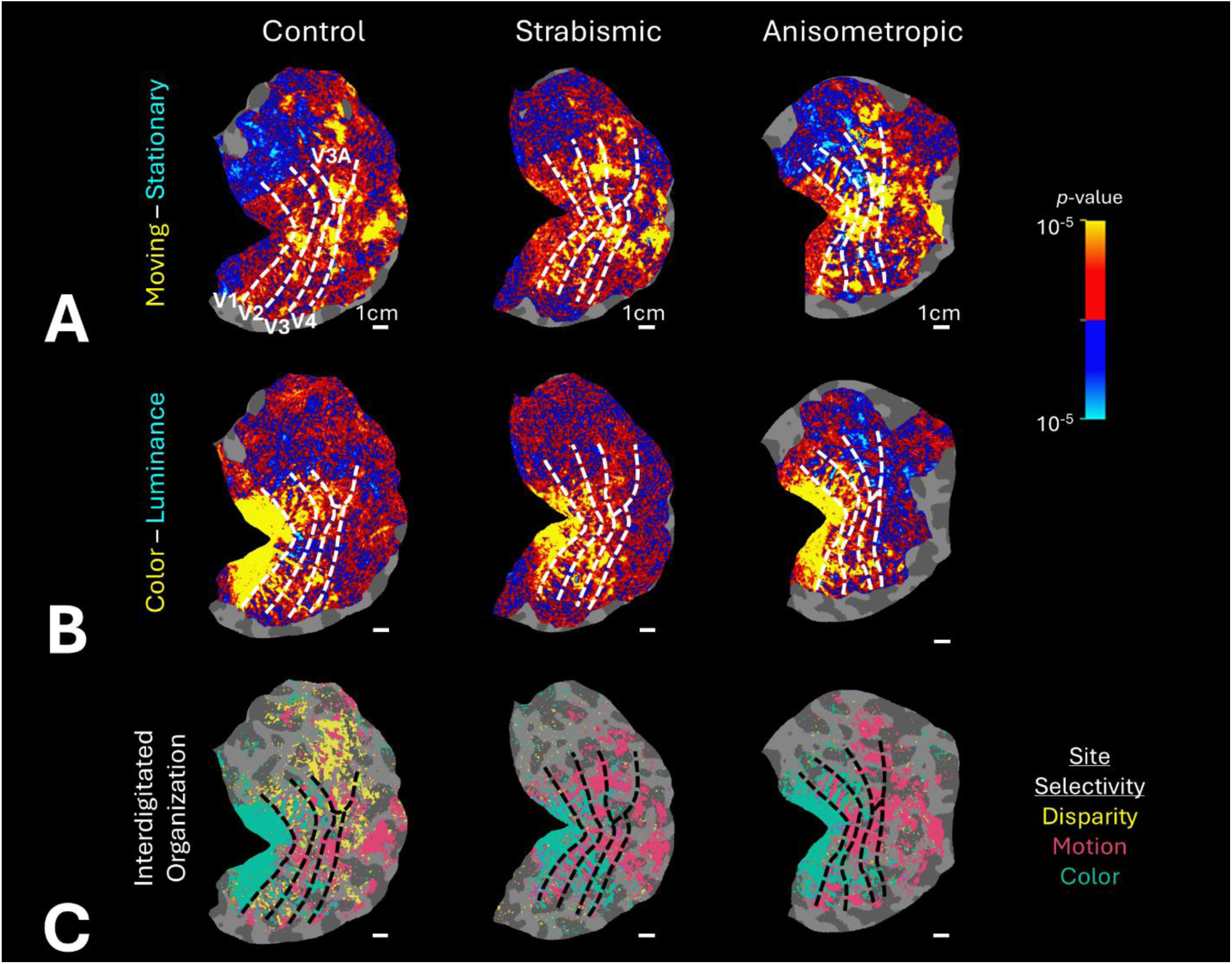
Interdigitation of disparity-, motion-, and color-selective activity in neurotypical and amblyopic vision. Panel **A** shows motion-selective activity, measured by contrasting responses to moving versus stationary rings. Panel **B** shows color-selective activity, measured by contrasting responses to color-varying versus achromatic luminance-varying stimuli. Panel **C** shows the corresponding interdigitation of sites exhibiting significant selective activity (*p* < 0.05; see Methods). Despite reduced disparity-selective activity in PwA, the interdigitation of motion- and color-selective sites remained evident. Other details are as in Figure 1.

When comparing the level of selective response across V2, V3, and V4 (Fig. 5A-B and Table S4), amblyopic participants exhibited significantly stronger activity within motion-preferring (F(1,29) = 4.41, *p* = 0.04) and color-preferring sites (F(1,29) = 4.72, *p* = 0.04) compared with controls, with no significant group × area interaction (F(2,58) = 0.99, *p* > 0.35). Whereas responses in stationary-preferring (Fig. 5C) and luminance-preferring sites (Fig. 5D) did not differ significantly between the amblyopic individuals and controls (F(1,29) < 1.71, *p* > 0.19) and there were also no group × area interactions (F(2,58) < 1.40, p > 0.26). However, this absence of group effect, specifically within the stationary preferring regions, appeared to be due to the difference between anisometropic and strabismic individuals (Fig. 5C) (see below).

**Figure 5).**
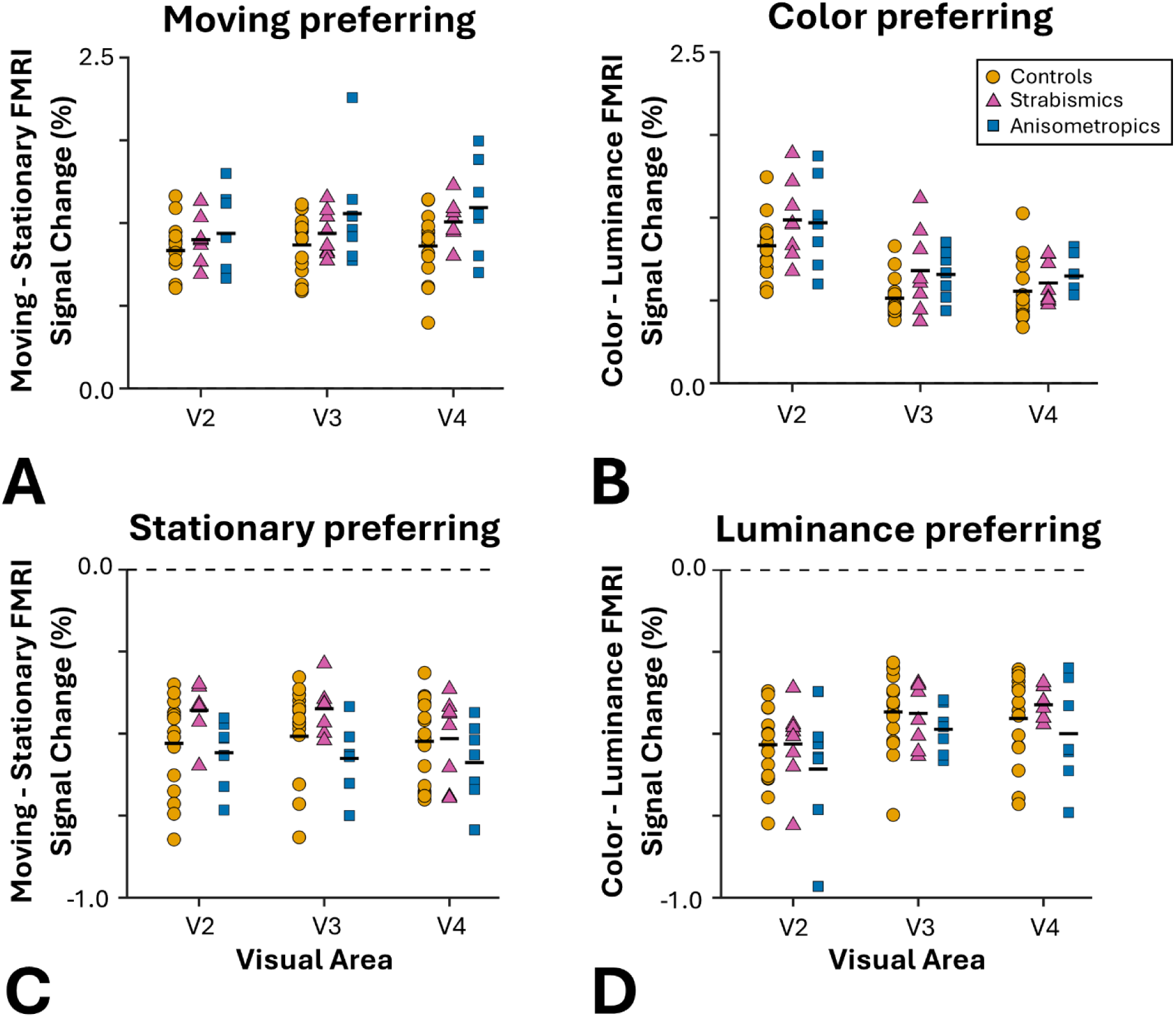
Selective activity within motion- vs. stationary-preferring and color- vs. luminance-preferring sites across V2–V4. Panels **A** and **B** show motion- and color-selective responses within motion-preferring and color-preferring sites, respectively. In both cases, PwA exhibited stronger selective activity than controls, with no apparent difference between strabismic and anisometropic individuals. Panels **C** and **D** show responses to the same stimuli within stationary-preferring and luminance-preferring sites. In stationary-preferring sites, strabismic individuals showed weaker (less negative) motion-selective activity compared with anisometropic individuals and controls. Other details are as in Figure 2.

To better clarify this phenomenon, we compared responses between strabismic and anisometropic participants. Activity in motion-preferring and color-preferring sites was comparable between the two amblyopia subtypes (F(1,29) < 0.60, *p* > 0.45) (Table S5). However, strabismic individuals showed significantly weaker (less negative) motion-selective responses in stationary-preferring sites (F(1,13) = 9.29, *p* < 0.01), while color-selective responses in luminance-preferring sites did not differ significantly between subtypes (F(1,13) = 1.76, *p* = 0.21). There were no significant group × area interactions (F(2,26) < 0.81, *p* > 0.42) (Table S5).

### 2.3. Expanded motion- and color-selective sites in amblyopia

We next asked whether the elevated motion- and color-selective responses in amblyopia were accompanied by an increase in the size of these regions. To test this, we compared the normalized size of regions that responded selectively to disparity, motion, and color (see Methods) across visual areas V2, V3, and V4, between controls and amblyopic individuals, adjusting for the total size of each visual area (Fig. 6).

**Figure 6).**
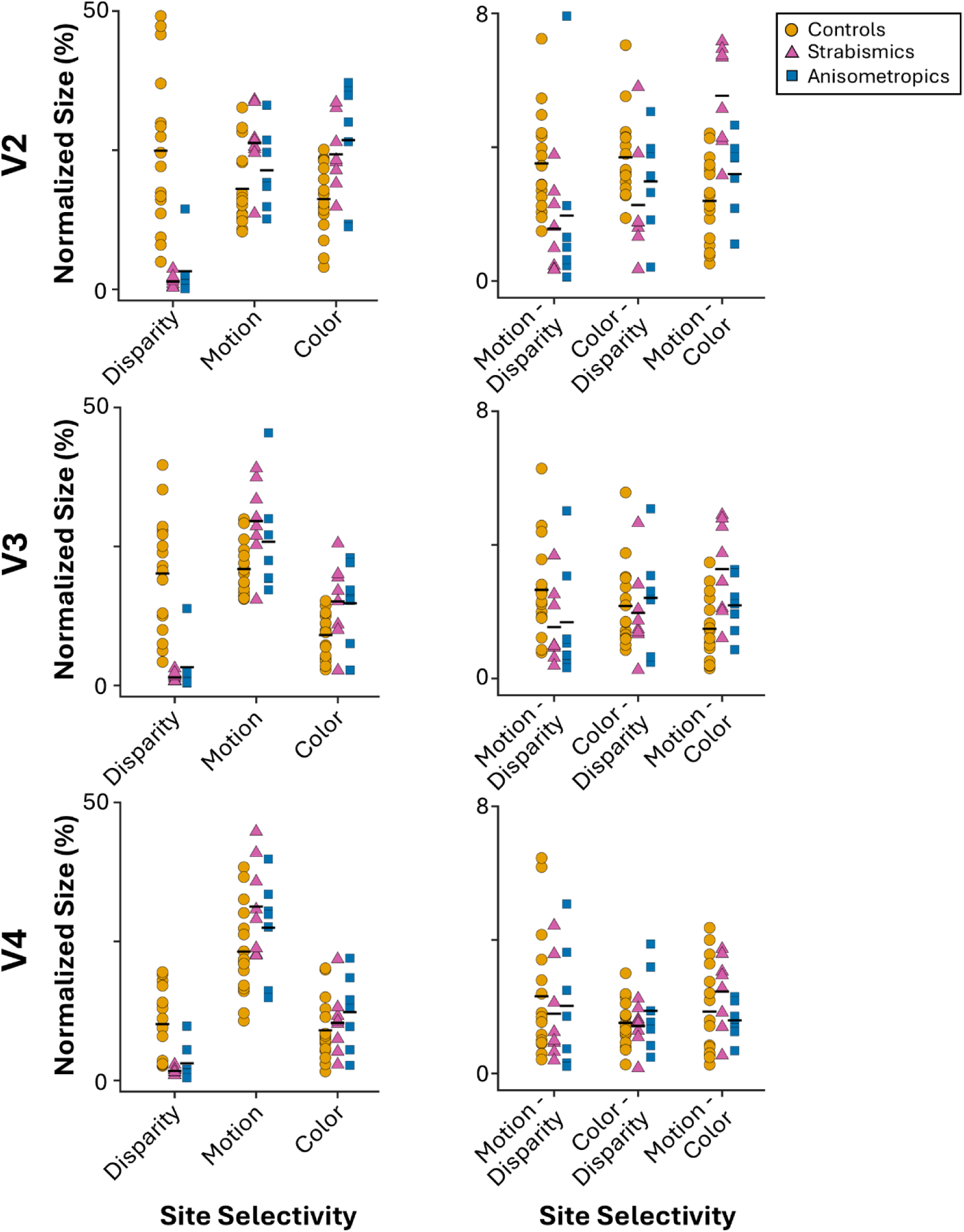
Size of disparity-, motion-, and color-selective sites and their level of overlap in controls and PwA. For each area, the top panel shows the size of non-overlapping selective sites, normalized to the size of the corresponding visual area. The bottom panels show the size of overlapping regions after normalization. In PwA, the reduction in the size of disparity-selective sites was accompanied by an increase in the size of motion- and color-selective sites. Strabismus, but not anisometropia, was further associated with increased overlap between motion- and color-selective sites.

In addition to a pronounced reduction in the size of disparity-preferring sites (F(1,29) = 36.64, p < 10⁻⁵), amblyopic individuals exhibited significantly larger motion- and color-selective regions compared with controls (F(1,29) = 14.01, *p* < 10⁻³), even when including sites showing overlapping selectivity with disparity-selective regions (Table S6). The size of motion- and color-selective sites remained comparable between subtypes (F(1,13) = 0.26, *p* = 0.61).

In our approach, the sizes of stereo-, motion-, and color-selective regions were calculated based on the relative magnitude of the evoked responses to these stimuli. One potential concern is that the smaller motion- and color-selective regions in controls compared to PwA could simply reflect the larger stereo-selective activity in these individuals. To address this, we re-estimated the sizes of the motion- and color-selective regions after nullifying the level of stereo-selective activity in both groups – i.e. independence in estimating the size of motion- and color-selective sites from the level of stereo-selective activity. The result of this test still revealed a significant main effect of group (F(1, 29) = 4.49, *p* = 0.04) due to larger motion- and color-selective sites in PwA compared to controls, with no interaction between the effect of group and the other independent parameters (*p* > 0.22). These findings indicate that the reduction in disparity-selective territory in amblyopia is accompanied by an expansion of motion- and color-selective regions. Importantly, this effect cannot be explained by methodological dependencies in our approach for estimating the relative sizes of motion- and color-selective versus stereo-selective regions.

### 2.4. Differential impact of amblyopia sub-types on the size of motion- and color-selective sites

The expansion in the size of motion- and color-selective sites was accompanied by a significant increase in the degree of overlap between them in PwA compared to controls (F(1,29) = 7.67, *p* < 0.01) even though this effect decreased significantly from V2 to V4 (area × group interaction; F(2, 58) = 14.26, *p* < 10^-4^). Importantly, this effect was significantly stronger in strabismic compared to anisometropic individuals (F(1,13) = 6.82, *p* = 0.02). This between sub-type difference decreased significantly from V2 to V4 (area × group interaction; F(2, 26) = 5.75, *p* = 0.02). Notably, while some degree of overlap between motion- and color-selective sites could reflect the spatial limits of fMRI and the partial voluming effect, these factors cannot explain the robust differences observed between the two amblyopia subtypes.

### 2.5. Evidence for developmental rivalry in controls

Our findings suggest that the reduction in disparity-selective activity in amblyopia is accompanied by an expansion of motion- and color-selective regions. However, correlations between these measures could not be assessed within the amblyopic group, because of the floor effect in disparity-selective responses and their representation. Instead, we tested for such relationships in controls.

As shown in Fig. 7, in V2, the size of disparity-selective sites was negatively correlated with the size of motion-selective sites (r = −0.65, *p* < 10⁻²) and color-selective sites (r = −0.79, *p* < 10⁻³). The negative correlation between disparity- and color-selective site size remained significant in V3 (r = −0.59, *p* = 0.02) and V4 (r = −0.56, *p* = 0.02), whereas the correlation between disparity- and motion-selective site size was not significant in these regions (*p* > 0.12). The size of motion- and color-selective sites was not significantly correlated in any of the three areas (*p* > 0.23). This result indicates that, in neurotypical individuals, the way motion- and color-selective responses compete with disparity-selective activity is likely additive and, rather independent from each other.

**Figure 7).**
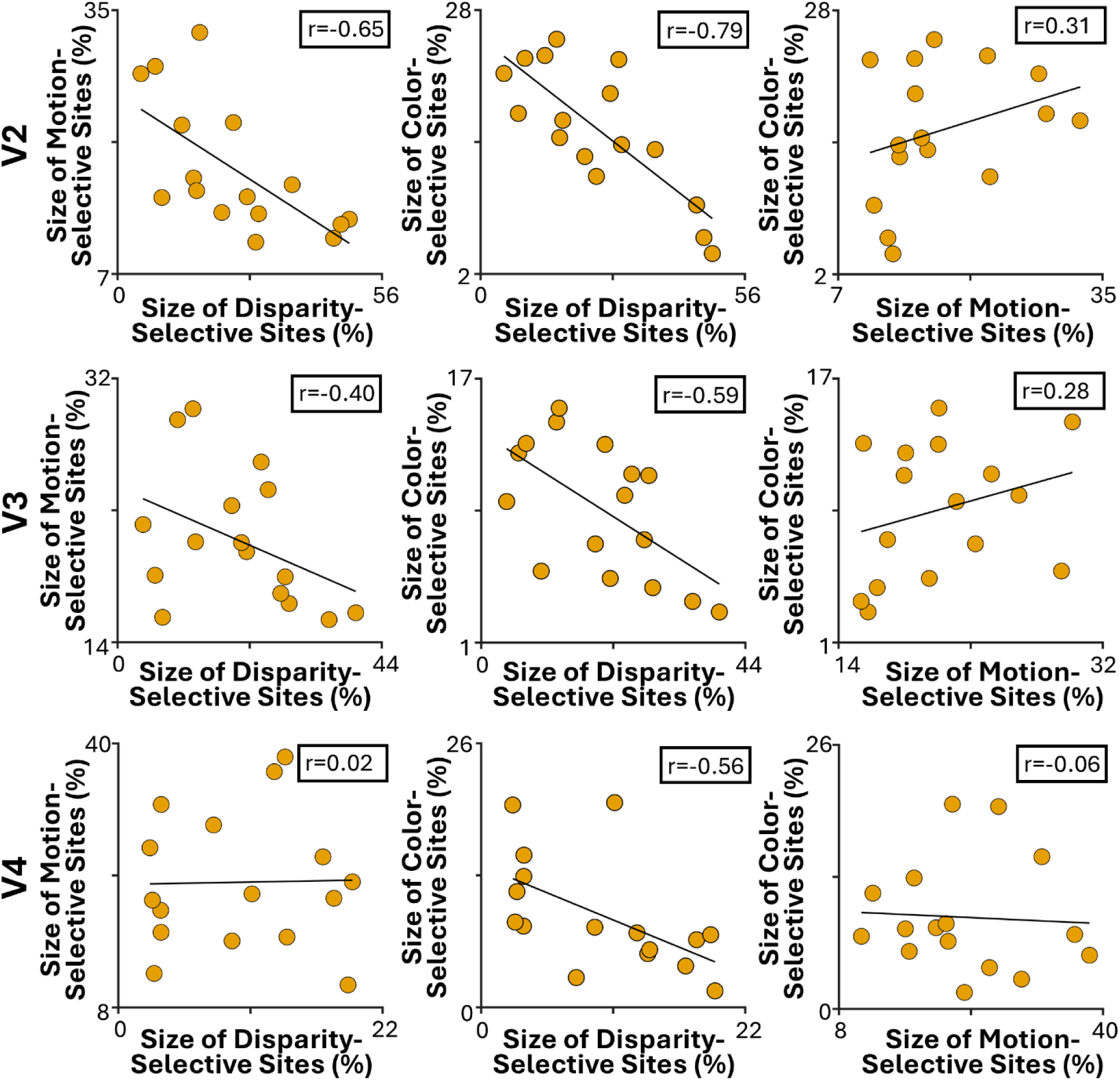
Correlation between the size of disparity-, motion-, and color-selective sites in neurotypical vision. The sizes of disparity-versus motion-selective sites and disparity-versus color-selective sites were negatively correlated. The strength of these correlations decreased from V2 to V4. No significant correlation was observed between the sizes of motion- and color-selective sites across V2–V4. Each data point represents one neurotypical participant.

## 3. Discussion

Amblyopia has long served as a primary model for studying visual cortical development, largely because of its well-documented impact on binocular functions such as depth perception (Levi, 2006; Levi et al., 2015). In this study, we provide evidence for decreased disparity-selective activity and demonstrate its correlation with amblyopia severity. Critically, our results also reveal that this disruption extends beyond disparity processing, influencing other cortical sites that support motion and color encoding; visual functions traditionally considered to be unaffected in PwA.

### 3.1. Developmental rivalry in individuals with amblyopic and neurotypical vision

Amblyopia is associated with a reduction in the number of neurons that respond preferentially to the amblyopic eye relative to the fellow eye (Crawford and von Noorden, 1979; Movshon et al., 1987; Smith et al., 1997; Kiorpes et al., 1998; Bi et al., 2011; Nasr et al., 2025). Our recent findings indicate that the magnitude of this interocular imbalance is correlated with amblyopia severity, indexed by the difference in visual acuity between the two eyes (Nasr et al., 2025). These results suggest a competitive interaction between the representations of the two eyes in primary visual cortex.

Here, we extend these findings to downstream visual areas, where the interocular imbalance is accompanied by a reduction in disparity-selective activity. Consistent with the behavioral findings (Reynaud and Hess, 2018; Wang et al., 2021), we showed that the extent to which disparity-selective responses were reduced correlated well with the difference in acuity between the two eyes, which gives a broad proxy of how binocular vision is compromised in amblyopia. Strikingly, we also find an expansion of motion- and color-selective sites, along with enhanced selective responses, providing one of the first demonstrations that amblyopia can be associated not only with selective loss of disparity-selective sites but also with reorganization of the other sites within the extrastriate cortex.

Furthermore, our findings revealed significant differences between the impact of amblyopia subtypes on the mesoscale functional organization of extrastriate visual areas. Compared to individuals with anisometropic amblyopia and controls, strabismic participants showed enlarged overlap between motion- and color-selective sites and reduced activity preference for stationary versus moving stimuli. This pattern suggests that strabismus more severely disrupts interactions between magnocellular and parvocellular pathways, thereby altering the balance between motion- and color-selective representations. These results go further in identifying differences between the impacts of strabismic vs. anisometropic amblyopia that, in the future, could be related to their behavioral measures.

Notably, evidence for developmental rivalry is not limited to PwA. In neurotypical individuals, we found that there was a negative correlation between the territory allocated to disparity vs. color and disparity vs. motion. We propose that this could be linked to the range of disparity sensitivity exhibited in control participants, which can be large.

### 3.2. The origin of developmental rivalry

Our results have shown that development can sculp the mesoscopic organization in extrastriate visual cortex. The nature of these changes prompts the question of where the changes may originate. Antecedent structures in the visual system, the lateral geniculate nucleus (LGN) and V1, are organized such that magno- and parvocellular streams are largely segregated. It is possible therefore that the signatures of amblyopia could be observed there and that we are sensitive to them in ‘reading out’ their effect in the extrastriate cortex, where, at the resolution of our experiments, we can visualize and quantify the mesoscopic organization related to the magnocellular and parvocellular streams.

However, to our knowledge, no study has reported an expansion of the color-selective (parvocellular) layers of LGN in amblyopic individuals. On the contrary, anatomical evidence indicates a reduction in the size of parvocellular neurons receiving input from the amblyopic eye (Hendrickson et al., 1987). Also, no previous study in NHPs assessed the impact of amblyopia on the organization of color- and motion-selective sites in V1. In the LGN, color-selective neurons are concentrated within the parvocellular layers. Considering this absence of knowledge, it would be valuable to investigate the layers of the LGN and color-selective blobs of V1, specifically to ask if amblyopia manifests in these structures.

### 3.3. Behavioral consequence of developmental rivalry

Whether the rivalry between the functional representation at mesoscopic scale has behavioral consequences remains to be seen. On the one hand, there has been the interesting observation that individuals with poor stereoacuity are overrepresented in art schools (Livingstone et al., 2011) consistent with the idea that higher perceptual skills associated with color processing may lend individuals an advantage in art. This observation is also consistent with some previous studies in PwA, showing increased sensitivity to stimuli presented to the fellow, but decreased sensitivity to the amblyopic, eye (Davis et al., 2006).

On the other hand, at least in PwA, the upregulation may just play a compensatory role that allows normal perception of color and motion. Specifically, in the absence of strong input to extrastriate visual areas from the amblyopic eye, more neurons are required for encoding visual features from a weaker visual signal. This alternative hypothesis aligns with the general consensus that, color and motion perception—particularly at higher contrast levels—are largely preserved in PwA, but so far, there is no behavioral evidence suggesting the superiority of PwA in color and motion tasks compared to controls.

### 3.4. Beyond amblyopia

Our findings raise questions about how congenital visual deficits of retinal origin may shape the mesoscopic functional organization of extrastriate cortex. For example, achromatopsia, a candidate for vision restoration therapies (Hoffmann and Dumoulin, 2015; Farahbakhsh et al., 2022), can result in exclusively rod-mediated vision during development. Reduced cone density has also been reported in individuals with albinism, while other congenital disorders such as Leber Congenital Amaurosis primarily affect rod development. Given the selective cortical processing of information encoded by these photoreceptors (e.g. see (Tootell and Nasr, 2021)), such disorders may have a lasting impact on the mesoscopic organization of the visual system and, in turn, influence the outcomes of restoration therapies. Thus, while previous studies have focused mainly on atrophic changes associated with developmental impairments such as amblyopia (Mendola et al., 2005; Liang et al., 2019) and achromatopsia (Lowndes et al., 2021; Molz et al., 2022), mesoscale functional changes in visual system organization should also be considered when evaluating impairment severity and predicting the potential outcomes of restoration therapies.

## 4. Conclusion

We introduce the concept of ‘developmental rivalry’ to describe competitive processes in visual development at the mesoscale levels. It manifests as an upregulation of the responses to and representation of functions that are spared at the expense of those which are reduced or lost. This result suggests that recruitment of brain real estate by other functions may serve as a barrier to recovery of stereo function. Our findings show that this rivalry unfolds differently in strabismic versus anisometropic amblyopia, with consequences likely extending beyond the measures reported here. This framework opens new directions for understanding the diverse manifestations of amblyopia.

## 5. Methods

### 5.1. Participants

Thirty-one human adults (14 females), aged 19–56 years old, participated in this study (Table S1). In this total of 31, there were 15 persons with amblyopia (7 with anisometropic, and 8 with strabismic amblyopia) and 16 individuals with normal or correct-to-normal visual acuity, who acted as controls. Outside the scanner, amblyopic participants were tested by an optometrist (J.S.) with extensive experience with amblyopic individuals to measure their best corrected visual acuity (ETDRS retro luminant chart (Precision Vision)). The stereoacuity was measured using Randot stereo test (Stereo Optical). We identified the participant’s dominant eye (Miles Test) and tested for suppression or diplopia (Worth 4 Dot).

All experimental procedures conformed to NIH guidelines and were approved by Massachusetts General Hospital protocols. Written informed consent was obtained from all participants prior to all experiments. The study conformed to the declaration of Helsinki and was approved by Massachusetts General Brigham (MGB) Institutional Review Board.

### 5.2. General Procedures

Participants were scanned multiple times in an ultra-high field 7T scanner (whole-body system, Siemens Healthcare, Erlangen, Germany) for functional experiments. All participants were also scanned in a 3T scanner (Tim Trio, Siemens Healthcare) for structural imaging.

In all functional experiments, stimuli were presented via an LCD projector (1024 × 768 pixel resolution, 60 Hz refresh rate) onto a rear-projection screen, viewed through a mirror mounted on the receive coil array. Matlab 2020a (MathWorks, Natick, MA, USA) and the Psychophysics Toolbox (Brainard, 1997; Pelli, 1997) were used to control stimulus presentation.

During the experiments, participants were instructed to look at a centrally presented object (radius = 0.15°) and to do an orthogonal dummy task (see below). These tasks were conducted irrespective of the stimuli presented in the background and remained the same across the whole run. In all experiments, the participant’s average performance for the dummy task remained above 75% without any significant difference across experimental conditions and/or between groups (*p*>0.10).

#### 5.2.1. Localizing disparity-selective columns

Disparity-selective sites were identified by comparing responses to disparity-varying stimuli (horizontal disparity: −0.22° to 0.22°; oscillated at 0.3 Hz) versus translationally moving, zero-disparity stimuli (left-to-right displacement: −0.22° to 0.22°; oscillates at 0.3 Hz). Stimuli consisted of sparse (5%) random dot stereograms (RDS) made of red or green dots (0.09° × 0.09°; 56 cd/m²), presented on a black background (Figure 1A) and spanning 20° × 26° of visual angle. Participants viewed the stimuli through custom-made anaglyphic spectacles integrated into the head coil. The experiment was block-designed with each block lasting 24 seconds, and each experimental run beginning and ending with 12 seconds of uniform black. Stimulus blocks were pseudo-randomized across runs to minimize order effects. Each participant completed two scanning sessions (168 s per run; 12 runs per session) on different days, during which the laterality of the anaglyphic lenses (red-left vs. red-right) was alternated and counterbalanced across participants. During the experiment, participants were instructed to look at a centrally presented object (radius = 0.15°). During the experiment, participants were asked to report shape-change for the fixation target (circle-to-square or vice versa).

#### 5.2.2. Localizing motion-selective columns

Motion-selective columns were localized based on their stronger response to moving compared to stationary stimuli. Stimuli consisted of concentric dark gray (30 cd/m^2^) rings, which were separated by 2 deg and had a width of 0.4deg, extending over 20° × 26° (height × width) in the visual field, and were presented against a light gray background (40 cd/m^2^). In half of the blocks, rings moved radially (centrifugally vs. centripetally) at 4°/s and the direction of motion changed every 3 s to reduce the impact of motion after-effects. In the other half of blocks, rings remained stationary during the whole block. The experiment was block-designed (24 s per block). Each run started and finished with a 12 s of uniform gray presentation (312 s per run; 6 runs per session). The sequence of moving and stationary blocks was pseudo-randomized across runs. During the experiments to localize motion and color-selective sites, participants were instructed to report color-change for the fixation target (red-to-blue or vice versa).

#### 5.2.3. Localizing color-selective columns

To localize color-selective columns, participants viewed sinusoidal gratings (0.2 cycles/deg; 20° × 26°) that varied either in chromaticity (alternating between red and blue) or in achromatic luminance. For each participant, red and blue hues were adjusted to be isoluminant across all stimulated eccentricities using the method of flicker photometry and tailored according to individual color perception (Nasr et al., 2016). The experiment was block-designed, with each stimulus block lasting 24 s. Each run started and finished with a 12 s of uniform gray presentation (312 s per run; 12 runs per session). The sequence of chromatic vs. luminance varying blocks was pseudo-randomized across runs.

During the experiments to localize motion and color-selective sites, participants were instructed to report color-change for the fixation target (dark-to-light green or vice versa).

#### 5.2.4. Retinotopic mapping

For all participants the border of retinotopic areas were defined by identifying representations of the vertical and horizontal meridians projected to surface reconstructions of the cortex (Sereno et al., 1995; Engel et al., 1997). Stimuli were pattern reversing (8 Hz) radial checkerboards, presented within limited apertures, against a gray background. These apertures limited the stimulus to wedge-shaped portions of the radial checkerboard orientated along the horizontal and vertical meridians with a width of 30° polar angle. The stimuli were presented to participants in different blocks (24 s per block). The sequence of blocks was pseudo-randomized across runs (8 blocks per run) and each participant was scanned in at least 4 runs (312 s per run).

### 5.3. Imaging

#### 5.3.1. Structural scans

We used anatomical images acquired at 3T on a Siemens TimTrio whole-body scanner, equipped with the standard vendor-supplied 32-channel RF head coil array, which provides more uniform image contrast over the full field of view (FOV) than imaging at 7T(Webb and Collins, 2010) (Wrede et al., 2012). We used a 3D T1-weighted MPRAGE sequence with protocol parameter values: TR=2530 ms, TE=3.39 ms, TI=1100 ms, flip angle=7°, bandwidth=200 Hz/pix, echo spacing=8.2 ms, voxel size=1.0×1.0×1.0 mm³, and FOV=256×256×170 mm³.

#### 5.3.2. Functional scans

We utilized a 7T whole-body Siemens scanner equipped with SC72 body gradients (70 mT/m gradient strength and max slew rate of 200 T/m/s), a custom-built 32-channel helmet radiofrequency (RF) receive coil array, and a birdcage volume RF transmit coil. Functional data were collected using single-shot 2D gradient-echo EPI, with 1.0 mm isotropic nominal voxel size, and the following protocol parameter values: TR=3000 ms, TE=28 ms, excitation flip angle=78°, FOV matrix=192×192, BW=1184 Hz/pixel, echo-spacing=1 ms, 7/8 phase partial Fourier, 44 oblique-coronal slices covering the occipital lobe, acceleration factor R=4 with GRAPPA reconstruction and FLEET-ACS data(Polimeni et al., 2015) with a 10° flip angle.

### 5.4. Data analysis

Functional and anatomical MRI data were pre-processed and analyzed using FreeSurfer (version 7.4.0; http://surfer.nmr.mgh.harvard.edu) (Fischl, 2012) and in-house developed MATLAB code. As described in the following sections in detail, each participant was analyzed entirely in their native anatomical space to preserve the fine-grained structure of cortical columns. This native-space processing was maintained throughout all individual analyses.

#### 5.4.1. Anatomical data

Based on each participant’s structural scan, cortical surfaces were reconstructed using FreeSurfer’s recon-all pipeline, which includes intensity normalization, skull stripping, white and pial surface extraction, and topological correction using a deformable surface model constrained by neuroanatomical priors(Dale et al., 1999). In this process, the standard pial surface was defined as the boundary between the gray matter (GM) and the surrounding cerebrospinal fluid (CSF), while the white matter (WM) surface was generated as the interface between the WM and GM. Additionally, nine intermediate surfaces were created using the equivolume approach, spaced at intervals of 10% of the cortical thickness. Finally, surfaces were flattened using FreeSurfer’s mris_flatten. To enhance the co-registration of functional and structural scans, all surfaces were upsampled to a 0.5 mm resolution using butterfly subdivision in mris_mesh_subdivide as implemented in FreeSurfer and outlined in our previously described pipeline (Wang et al., 2022).

#### 5.4.2. Functional data

The collected functional data were first upsampled to 0.5 mm isotropic resolution using trilinear interpolation as implemented in FreeSurfer’s mri_convert (see also(Wang et al., 2022)). For each participant, the functional data from each run were rigidly aligned (with 6 degrees of freedom) to their structural scan using rigid Boundary-Based Registration (Greve and Fischl, 2009). Head motion covariates were then derived for six directions. To preserve spatial resolution on the surface, no tangential spatial smoothing was applied to the fMRI data. Instead, functional data was projected to the 11 cortical surfaces of each individual and radial (intracortical) smoothing— perpendicular to the cortex and within cortical columns—was used(Blazejewska et al., 2019). To minimize the blurring impact of pial veins on functional map resolution(Koopmans et al., 2010; Polimeni et al., 2010; De Martino et al., 2013; Nasr et al., 2016), radial smoothing was restricted to the deep cortical depth (as defined in 2.4.1). Notably, we avoided applying any method of spatial distortion correction to the fMRI data because EPI distortions are typically small in the occipital lobe (Renvall et al., 2016).

After preprocessing, a standard hemodynamic response model based on a gamma function was fitted to the signals from deep cortical depths to estimate the amplitude of the blood-oxygen-level-dependent (BOLD) response. To generate disparity-, motion-, and color-selectivity maps, a vertex-wise general linear model was applied by computing the contrast of interest (Nasr et al., 2016, 2025).

### 5.5. Region of interest (ROI) analysis

#### 5.5.1. Amplitude of the evoked response across visual areas

To measure the amplitude of disparity-, motion-, and color-selective activity across visual areas, we first divided each area into sites preferring one of the rival stimulus types (e.g., moving vs. stationary stimuli), based on the unthresholded maps. The level of selective activity was then measured separately within each site by subtracting the response evoked by each of the rival stimuli; e.g. motion-selective response was calculated as the difference between the response to moving vs. stationary stimuli. This approach prevented cancellation effects that would arise from averaging across the entire visual area.

#### 5.5.2. Estimating the size of mesoscale regions and analysis of overlap

Our overall approach is similar to those described previously elsewhere (Kennedy et al., 2023). Briefly, to measure the size of disparity-, motion, and color-selective sites and then the level of overlap between sites, the selectivity maps (Figures 1 and 4) were first thresholded at *p*<0.05. This step assured us that we were not arbitrarily assigning a label (e.g. disparity-selective) to a site due to noise in our measurements. Then, the level of activity across these thresholded sites were normalized (min-to-max were converted linearly to 0-to-1) after excluding the outlier values. A cortical site was called “selective” if the resultant normalized value for one feature (i.e. either disparity, motion, or color) was 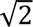 time larger than that of the value measured for any of the others. A cortical size was considered “overlap” when the ratio of normalized values for two features was smaller than 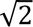.

### 5.6. Statistical analysis

Repeated measured ANOVA was used primarily to test the impact of two planned contrasts; 1) amblyopic vs. control participants and 2) strabismic vs. anisometropic participants. Since this analysis is particularly susceptible to the violation of sphericity assumption, caused by the correlation between measured values, when necessary (determined using a Mauchly test), results were corrected for violation of the sphericity assumption, using the Greenhouse-Geisser method. All post-hoc analyses were conducted after Bonferroni correction for multiple comparisons.

### 5.7. Data availability statement

Data and codes will be shared upon request.

## Acknowledgment

This work was supported by NIH NEI (R01 EY030434 and R01 EY017081), and by the MGH/HST Athinoula A. Martinos Center for Biomedical Imaging, Max Planck School of Cognition, the German Federal Ministry of Education and Research (BMBF) and the Max Planck Society. Crucial resources were made available by a NIH Shared Instrumentation Grant S10-RR019371. We thank Ms. Azma Mareyam for helping with hardware maintenance during this study. We thank Ms. Amanda Nabasaliza for her help with the recuitment.

