## Supplementary Tables for "Mesoscale developmental rivalry in human extrastriate visual cortex"

**Table S1.** Demography

| *ID* | *Gender* | *Age* | *Age at Diagnosis* | *Right Eye Visual Acuity* | *Left Eye Visual Acuity* | *Binocular Visual Acuity* | *Fellow / Dominant Eye* | *Suppression Worth 4Dots* | *Randot Stereoacuity* | *Angle of Strabismus (at 4 m in PD)* |
|---|---|---|---|---|---|---|---|---|---|---|
| S1 | F | 40 | <1 | -0.06 | +0.08 | -0.06 | RE | Diplopia | >500 | 16/25 |
| S2 | M | 19 | 3 | +0.09 | +0.30 | +0.09 | LE | RE | >500 | 12/10 |
| S3 | M | 28 | 6 | +0.48 | -0.02 | +0.02 | LE | RE | >500 | 25/18 |
| S4 | F | 31 | 3 | +0.26 | +0.04 | +0.04 | LE | RE | >500 | 10/8 |
| S5 | F | 26 | 2 | +0.46 | -0.06 | -0.14 | LE | RE | >500 | 10/8 |
| S6 | F | 20 | 4 | -0.08 | -0.10 | -0.08 | LE | Diplopia | 70 | 20/20 |
| S7 | M | 56 | 5 | -0.20 | +0.06 | +0.06 | RE | LE | >500 | 16/16 |
| S8 | M | 28 | 2 | +0.00 | +0.60 | +0.00 | RE | LE | >500 | 14/10 |
| A1 | F | 31 | 5 | -0.22 | +0.20 | -0.16 | RE | None | >500 | None |
| A2 | M | 23 | 5 | -0.08 | +0.30 | -0.04 | RE | None | 400 | None |
| A3 | M | 35 | 11 | -0.04 | +0.26 | -0.04 | RE | LE | 200 | None |
| A4 | M | 26 | 8 | +0.00 | +0.17 | +0.00 | RE | Diplopia | 200 | None |
| A5 | F | 19 | 5 | +1.00 | -0.08 | -0.08 | LE | RE | >500 | None |
| A6 | F | 24 | 8 | +0.64 | -0.10 | -0.10 | LE | RE | >500 | None |
| A7 | M | 29 | 4 | -0.26 | +0.10 | -0.26 | RE | LE | 100 | None |

**Table S2.** Group: Controls (n=16) vs. Amblyopes (n=15); Site: Disparity preferring vertices; Area: V2 vs. V3 vs. V4

| ***Effect*** | ***df*** | ***F*** | ***p*** |
|---|---|---|---|
| ***Group*** | 1, 29 | 16.57 | $<10^{-3}$*** |
| ***Area*** | 2, 58 | 4.03 | 0.048* |
| ***Group* × *Area*** | 2, 58 | 20.94 | $<10^{-4}$*** |

Group: Controls (n=16) vs. Amblyopes (n=15); Site: Zero-disparity preferring vertices; Area: V2 vs. V3 vs. V4

| ***Effect*** | ***df*** | ***F*** | ***p*** |
|---|---|---|---|
| ***Group*** | 1, 29 | 0.06 | 0.81 |
| ***Area*** | 2, 58 | 8.43 | $<10^{-3}$*** |
| ***Group* × *Area*** | 2, 58 | 2.94 | 0.07 |

df = degrees of freedom; *: $p<0.05$; **: $p<0.01$; ***: $p<10^{-3}$

**Table S3.** Group: Strabismics (n=8) vs. Anisometropes (n=7); Site: Disparity preferring vertices; Area: V2 vs. V3 vs. V4

| ***Effect*** | ***df*** | ***F*** | ***p*** |
|---|---|---|---|
| ***Group*** | 1, 13 | 0.40 | 0.54 |
| ***Area*** | 2, 26 | 9.39 | <0.01** |
| ***Group* × *Area*** | 2, 26 | 1.77 | 0.20 |

Group: Strabismics (n=8) vs. Anisometropes (n=7); Site: Zero-disparity preferring vertices; Area: V2 vs. V3 vs. V4

| ***Effect*** | ***df*** | ***F*** | ***p*** |
|---|---|---|---|
| ***Group*** | 1, 13 | 0.01 | 0.92 |
| ***Area*** | 2, 26 | 9.46 | <0.01** |
| ***Group* × *Area*** | 2, 26 | 0.56 | 0.55 |

df = degrees of freedom; **: $p<0.01$

**Table S4.** Group: Controls (n=16) vs. Amblyopes (n=15); Site: Motion-only; Area: V2 vs. V3 vs. V4
Motion preferring:

| *Effect* | *df* | *F* | *p* |
|---|---|---|---|
| ***Group*** | 1, 29 | 4.41 | 0.04* |
| ***Area*** | 2, 58 | 3.79 | 0.03* |
| ***Group* × *Area*** | 2, 58 | 1.57 | 0.22 |

Static preferring:

| *Effect* | *df* | *F* | *p* |
|---|---|---|---|
| ***Group*** | 1, 29 | 0.05 | 0.82 |
| ***Area*** | 2, 58 | 1.74 | 0.19 |
| ***Group* × *Area*** | 2, 58 | 1.40 | 0.26 |

Group: Controls (n=16) vs. Amblyopes (n=15); Site: Color-only; Area: V2 vs. V3 vs. V4

Color preferring:

| *Effect* | *df* | *F* | *p* |
|---|---|---|---|
| ***Group*** | 1, 29 | 4.72 | 0.04* |
| ***Area*** | 2, 58 | 96.89 | $<10^{-15}$*** |
| ***Group* × *Area*** | 2, 58 | 1.71 | 0.19 |

Luminance preferring:

| *Effect* | *df* | *F* | *p* |
|---|---|---|---|
| ***Group*** | 1, 29 | 0.26 | 0.61 |
| ***Area*** | 2, 58 | 20.12 | $<10^{-5}$*** |
| ***Group* × *Area*** | 2, 58 | 0.49 | 0.57 |

df = degrees of freedom; *: $p<0.05$; **: $p<0.01$; ***: $p<10^{-3}$

**Table S5.** Group: Strabismics (n=8) vs. Anisometropes (n=7); Site: Motion-only; Area: V2 vs. V3 vs. V4

Motion preferring:

| *Effect* | *df* | *F* | *p* |
|---|---|---|---|
| ***Group*** | 1, 13 | 0.60 | 0.45 |
| ***Area*** | 2, 26 | 5.00 | 0.02* |
| ***Group* × *Area*** | 2, 26 | 0.47 | 0.59 |

Static preferring:

| *Effect* | *df* | *F* | *p* |
|---|---|---|---|
| ***Group*** | 1, 13 | 9.29 | <0.01** |
| ***Area*** | 2, 26 | 1.97 | 0.17 |
| ***Group* × *Area*** | 2, 26 | 0.81 | 0.42 |

Group: Strabismics (n=8) vs. Anisometropes (n=7); Site: Color-only; Area: V2 vs. V3 vs. V4

Color preferring:

| *Effect* | *df* | *F* | *p* |
|---|---|---|---|
| ***Group*** | 1, 13 | 0.00 | 0.99 |
| ***Area*** | 2, 26 | 42.13 | $<10^{-6}$*** |
| ***Group* × *Area*** | 2, 26 | 0.38 | 0.64 |

Luminance preferring:

| *Effect* | *df* | *F* | *p* |
|---|---|---|---|
| ***Group*** | 1, 13 | 1.76 | 0.21 |
| ***Area*** | 2, 26 | 9.51 | <0.01** |
| ***Group* × *Area*** | 2, 26 | 0.24 | 0.75 |

df = degrees of freedom; *: $p<0.05$; **: $p<0.01$; ***: $p<10^{-3}$

**Table S6.** Group: Controls (n=16) vs. Amblyopia (n=15); Site Type: Color vs. Motion; Area: V2 vs. V3 vs. V4

Non-overlapping parts:

| *Effect* | *df* | *F* | *p* |
|---|---|---|---|
| ***Group*** | 1, 29 | 14.01 | $<10^{-3}$*** |
| ***Type*** | 1, 29 | 20.51 | $<10^{-4}$*** |
| ***Area*** | 2, 58 | 28.99 | $<10^{-8}$*** |
| ***Group × Type*** | 1, 29 | 0.00 | 0.97 |
| ***Group × Area*** | 2, 58 | 2.01 | 0.15 |
| ***Type × Area*** | 2, 58 | 2.15 | 0.14 |

Overlapping parts:

| *Effect* | *df* | *F* | *p* |
|---|---|---|---|
| ***Group*** | 1, 29 | 7.67 | <0.01** |
| ***Area*** | 2, 58 | 50.95 | $<10^{-10}$*** |
| ***Group × Area*** | 2, 58 | 14.26 | $<10^{-4}$*** |

Group: Strabismics (n=8) vs. Anisometropes (n=7); Site Type: Color vs. Motion; Area: V2 vs. V3 vs. V4

Non-overlapping parts:

| *Effect* | *df* | *F* | *p* |
|---|---|---|---|
| ***Group*** | 1, 13 | 0.26 | 0.62 |
| ***Type*** | 1, 13 | 6.51 | 0.02* |
| ***Area*** | 2, 26 | 31.82 | $<10^{-6}$*** |
| ***Group × Type*** | 1, 13 | 0.64 | 0.44 |
| ***Group × Area*** | 2, 26 | 0.53 | 0.59 |
| ***Type × Area*** | 2, 26 | 0.27 | 0.69 |

Overlapping parts:

| *Effect* | *df* | *F* | *p* |
|---|---|---|---|
| ***Group*** | 1, 13 | 6.82 | 0.02* |
| ***Area*** | 2, 26 | 52.38 | $<10^{-6}$*** |
| ***Group × Area*** | 2, 26 | 5.75 | 0.02* |

df = degrees of freedom; *: $p<0.05$; **: $p<0.01$; ***: $p<10^{-3}$
